# Lipoprotein Signatures of Cholesteryl Ester Transfer Protein and HMG-CoA Reductase Inhibition

**DOI:** 10.1101/295394

**Authors:** Johannes Kettunen, Michael V. Holmes, Elias Allara, Olga Anufrieva, Pauli Ohukainen, Clare Oliver-Williams, Therese Tillin, Alun D. Hughes, Mika Kähönen, Terho Lehtimäki, Jorma Viikari, Olli T. Raitakari, Veikko Salomaa, Marjo-Riitta Järvelin, Markus Perola, George Davey Smith, Nish Chaturvedi, John Danesh, Emanuele Di Angelantonio, Adam S. Butterworth, Mika AlaKorpela

## Abstract

**Background:** CETP inhibition reduces vascular event rates but confusion surrounds its low-density lipoprotein (LDL)-cholesterol effects. We sought to clarify associations of genetic inhibition of CETP on detailed lipoproteins.

**Methods and Results:** We used variants associated with *CETP* (rs247617) and *HMGCR* (rs12916) expression in 62,400 Europeans with detailed lipoprotein profiling from nuclear magnetic resonance spectroscopy. Genetic associations were scaled to 10% lower risk of coronary heart disease (CHD). Associations of lipoprotein measures with risk of incident CHD in three population-based cohorts (770 cases) were examined.

*CETP* and *HMGCR* had near-identical associations with LDL-cholesterol concentration estimated by Friedewald-equation. *HMGCR* had a relatively consistent effect on cholesterol concentrations across all apolipoprotein B-containing lipoproteins. *CETP* had stronger effects on remnant and very-low-density lipoprotein cholesterol but no effect on cholesterol concentrations in LDL defined by particle size (diameter 18–26 nm) (-0.02SD 95%CI: -0.10, 0.05 for *CETP* versus -0.24SD, 95%CI -0.30, -0.18 for *HMGCR*). *CETP* had profound effects on lipid compositions of lipoproteins, with strong reductions in the triglyceride content of all highdensity lipoprotein (HDL) particles. These alterations in triglyceride composition within HDL subclasses were observationally associated with risk of CHD, independently of total cholesterol and triglycerides (strongest HR per 1-SD higher triglyceride composition in very-large HDL 1.35; 95%CI: 1.18, 1.54).

**Conclusion:** CETP inhibition does not affect size-specific LDL cholesterol but may lower CHD risk by lowering cholesterol in other apolipoprotein-B containing lipoproteins and lowering triglyceride content of HDL particles. Conventional composite lipid assays may mask heterogeneous effects of lipid-altering therapies.

## Introduction

Definitive evidence on the causal role of low-density lipoproteins (LDL) in cardiovascular disease comes from trials of LDL cholesterol lowering compounds,^1^ which have shown beneficial effects on risk of coronary heart disease (CHD) and stroke. Consistent effects have been seen for drugs acting on related pathways, such as 3-hydroxy-3-methylglutaryl-coenzyme A reductase (HMGCR) inhibitors, i.e., statins, and proprotein convertase subtilisin-kexin type 9 (PCSK9) inhibitors,^2^ both of which upregulate hepatic LDL receptor expression, and for drugs acting on other pathways, such as ezetimibe, which inhibits intestinal absorption of cholesterol.^3^

However, trials of drugs primarily designed to alter concentrations of lipids other than LDL cholesterol have had mixed results.^4^ One such example is the class of drugs designed to inhibit cholesteryl ester transfer protein (CETP), a lipid transport protein responsible for the exchange of triglycerides and cholesteryl esters between apolipoprotein B-containing particles and high-density lipoprotein (HDL) particles. CETP inhibitors were developed initially on the basis of their HDL cholesterol raising effects. While accumulating genetic evidence suggests that HDL cholesterol concentration is unlikely to be causally related to CHD,^5^ there were two strong reasons to believe that CETP inhibition may still reduce vascular risk: (i) genetic studies of *CETP* variants have shown associations with CHD^6^ and (ii) some CETP inhibitors not only increase HDL cholesterol but also appear to lower LDL cholesterol as measured by conventional assays.^7^

The recent findings from the phase III REVEAL trial showed that treatment with the CETP inhibitor anacetrapib led to a reduction in risk of coronary events that was proportional to the reduction in non-HDL cholesterol.^8^ Interestingly, anacetrapib appeared to have discrepant effects based on the assay used to quantify LDL cholesterol (using beta-quant, direct or Friedewald estimation).^7^ This discrepant effect was also identified in a genetic study that approximated a factorial clinical trial of CETP inhibition and statin therapy.^9^ Thus, while both CETP inhibitors and statins lower Friedewald-estimated LDL cholesterol, which also includes cholesterol carried by other lipoprotein particles, it is possible that the drugs have differential effects on the concentration and content of lipids in different apolipoprotein-B containing lipoproteins.

In this study, we used the established approach of exploiting genetic variants near the protein-coding genes of drug targets to investigate detailed lipid and lipoprotein subclass signatures of CETP inhibition. We compared the association of variants in *CETP* with *HMGCR*^10^ (to proxy statin treatment) to gauge insight into how these two therapies alter the lipoprotein milieu. We also present findings that the triglyceride composition, in contrast to circulating concentrations, of HDL particles is associated with CHD and may relate to a new mechanism by which CETP inhibition reduces risk of CHD.

## Methods

### Prospective and cross-sectional studies and lipoprotein quantification

We used genetic and lipoprotein data from four population-based Finnish cohorts and one cross-sectional study in the UK (cohort characteristics are presented in Online Table 1 and study descriptions are given in the Supplementary Note in the Supplementary Appendix). For prospective analyses we used two of the abovementioned Finnish cohorts and additionally a UK-based multiethnic SABRE (Southall And Brent REvisited) cohort. Briefly, the cohorts used were the Northern Finland Birth Cohort 1966 (NFBC66) (*n* = 4,702 individuals aged 31 y at blood draw),^11^ the Cardiovascular Risk in Young Finns Study (YFS, *n* = 1,948 individuals aged 24–39 y in 2007),^12^ two population based Finnish cohorts FINRISK 1997 (*n* = 6,942 individuals aged 24–74 y) and DILGOM subsample of FINRISK 2007 (*n* = 4,124 individuals aged 24–74 y),^13^ a study of healthy blood donors from the UK (INTERVAL: *n* = 40,958 individuals aged 18–80 y)^14^ and a tri-ethnic UK community-based cohort SABRE (*n* = 4,857 individuals aged 40–69 y).^15^ A nuclear magnetic resonance (NMR)-based methodology was used to quantify lipoprotein lipids and subclasses. Details of this platform have been published previously,^16, 17^ and it has been widely applied in genetic and epidemiological studies.^10, 18^

Where possible, we excluded individuals receiving lipid lowering medication, pregnant women and those who had a high proportion (>30%) of values missing across the lipid traits. All measures were first adjusted for sex, age (if applicable), genotyping batch (if applicable) and ten first principal components from genomic data and the resulting residuals were transformed to normal distribution by inverse rank-based normal transformation. Details of study-specific genotyping are provided in Online Table 2 in the Supplementary Appendix.

### SNP analysis

We selected variants as genetic proxies of CETP and HMGCR inhibition on the basis of robust associations with circulating lipids in GWAS consortia^19, 20^ and target gene expression. The *HMGCR* variant (rs12916) LDL cholesterol lowering T allele (-0.24 SD LDL cholesterol per T allele; P=1.3×10^−14^) has been shown to lower *HMGCR* expression^21^ and the *CETP* variant (rs247617) HDL cholesterol increasing A allele (0.84 SD HDL cholesterol per A allele; P=5.4×10^−94^) associates with lower gene expression across several tissues, verified through Genotype To Expression (https://gtexportal.org) project data.^22^ We used an additive model for each cohort separately (see Online Table 1 for details of analysis software). In order to make the lipoprotein and lipid estimates comparable, the estimates for *CETP* rs247617 and *HMGCR* rs12916 were scaled to the same CHD association as reported by the CARDIoGRAMplusC4D GWAS Consortium.^23^ The per-allele log odds (logOR) for CHD was 0.0358 for *HMGCR* rs12916 and 0.0309 for *CETP* rs247617; subsequently the summary statistics of each individual cohort and each metabolite were scaled to -0.105 logOR of CHD (equivalent to an odds ratio [OR] of CHD of 0.90) to align the estimates to a 10% lower risk of CHD. The cohort specific association results of lipoprotein and lipid measures with both variants were then combined using an inverse variance weighted fixed effect meta-analysis.

Our focus for this study was to evaluate the impact of variants in *CETP* and *HMGCR* on the entire cascade of apolipoprotein B-containing lipoproteins and HDL subclasses. Therefore, we decided *a priori* to examine 191 lipoprotein and lipid traits available from the NMR platform.^18, 24^ Focusing on these 191 traits, we estimated that 28 principal components explain 99% of their variation in the Finnish cohorts and therefore we used a P-value threshold of 0.05/28=0.002 to denote evidence in favor of an association. Abbreviations and full descriptions of the lipoprotein measures studied are listed in Online Table 3.

### Association of lipoprotein measures with risk of incident CHD

Cohorts contributing to the associations of lipoprotein lipid concentration and composition measures and the hazard of incident CHD were FINRISK 1997, DILGOM and SABRE. Participants with prevalent CHD were excluded from the analysis. Following exclusion, data were available from FINRISK 1997 for 7,076 individuals (291 cases / 6,785 controls) and 4,736 individuals from DILGOM (192 cases / 4,544 controls) and for SABRE 4,689 individuals with non-missing data (287 cases / 4,402 controls). The follow up time of FINRISK 1997 and SABRE were censored to 8 years to match the follow up time in DILGOM.

Prior to statistical analyses, metabolic measures were log-transformed and scaled to standard deviations (SD) in each cohort. The relationships of lipid measures with the risk of CHD were analysed using Cox proportional hazards regression models with age, sex, mean arterial pressure, smoking, diabetes mellitus, lipid medication and geographical region (Finnish cohorts), ethnicity (SABRE), total cholesterol and total triglyceride concentrations as covariates. The cohort-specific association results of 191 lipid measures were then combined using an inverse variance weighted fixed effect meta-analyses. Analyses were conducted in R studio (version 1.0.153, R version 3.3.3). As above, we used a P-value threshold of ≤0.002 to denote evidence in favor of an association.

## Results

Data from 62,400 individuals with extensive lipoprotein subclass profiling and genotypes were available. We combined data from five adult cohorts (mean age range from 31 to 52 years) and one cohort of adolescents (mean age 16 years) for the genetic analyses where 51% of participants of all six studies were female. Study specific and pooled estimates from meta-analyses of genetic and observational analyses for all 191 traits are presented in Online Figures 1-15.

**Figure 1.**
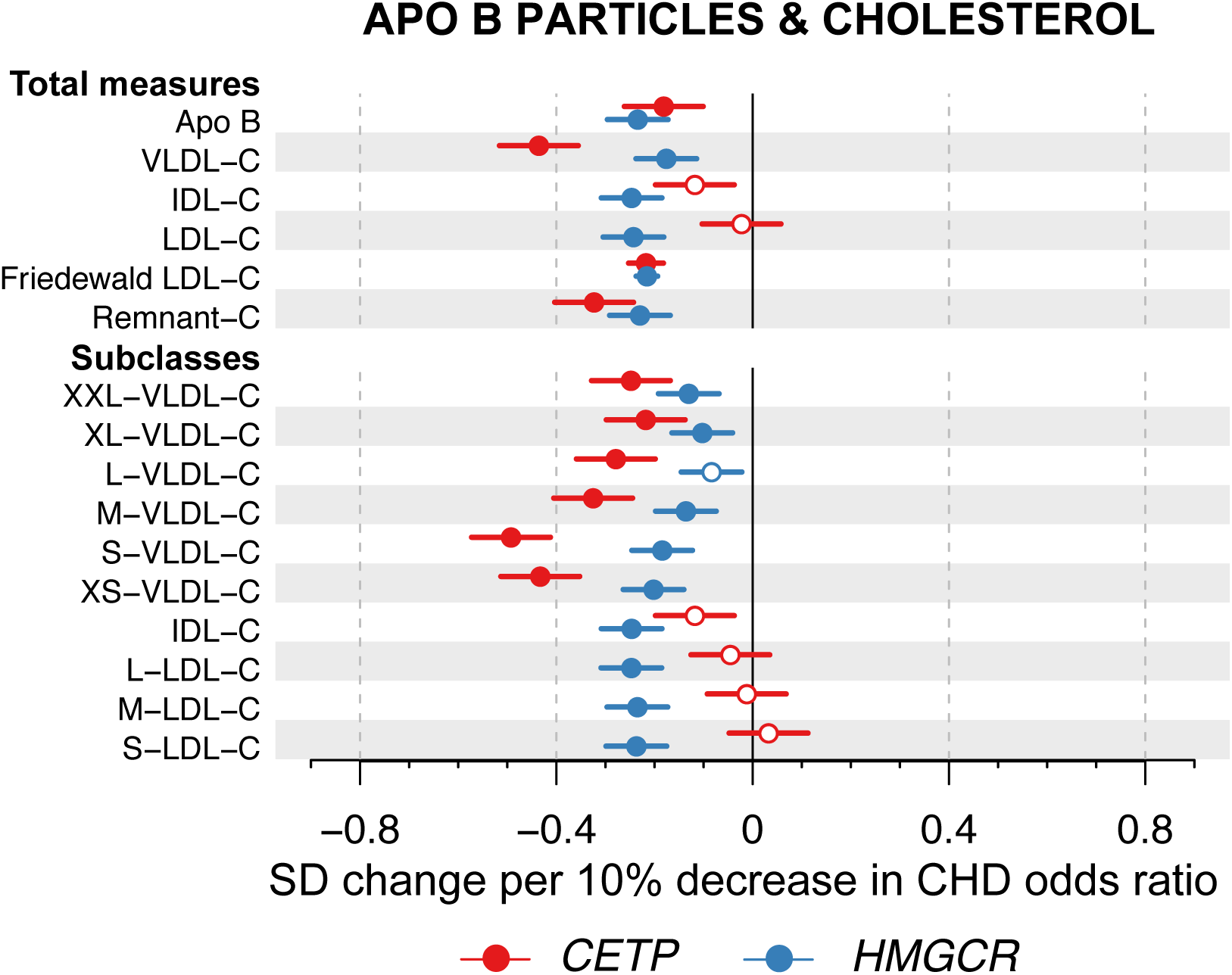
Associations of genetic variants in *CETP* rs247617 (red) and *HMGCR* rs12916 (blue) with circulating apolipoprotein B and cholesterol concentrations in size-specific apolipoprotein B particles. Estimates represent the standardized difference in lipoprotein trait, with per-allele associations scaled to a 10% lower risk of CHD. Analyses were adjusted for age, sex, genotyping batch and ten genetic principal components. Closed circles represent statistical significance of associations at P<0.002 and open circles associations that are non-significant at this threshold. The lipoprotein subclasses are defined by particle size:^17, 18, 25^ potential chylomicrons and the largest very-low-density lipoprotein particles (XXL-VLDL; average particle diameter ≥75 nm); five different VLDL subclasses, i.e. very large (average particle diameter 64.0 nm), large (53.6 nm), medium (44.5 nm), small (36.8 nm) and very small VLDL (31.3 nm); intermediate-density lipoprotein (IDL; 28.6 nm); and three LDL subclasses, i.e. large (25.5 nm), medium (23.0 nm) and small LDL (18.7 nm).

Scaled to a 10% lower risk of CHD, *CETP* rs247617 and *HMGCR* rs12916 had near-identical associations with Friedewald estimated LDL cholesterol (Fig. 1) and similar associations for apolipoprotein B. In contrast, when LDL cholesterol was defined on the basis of cholesterol transported in LDL based on particle size (diameter 18–26 nm), and measured via NMR spectroscopy, *CETP* had no association with this size-specific LDL cholesterol (0.02 SDs; 95%CI: -0.10, 0.05). While *HMGCR* had a relatively consistent association with individual apolipoprotein B-containing lipoproteins (effect estimates ranging from -0.25 for IDL cholesterol to -0.18 for VLDL cholesterol), *CETP* had the most pronounced associations with VLDL cholesterol, a weaker association with IDL cholesterol but no association with LDL cholesterol defined by particle size or cholesterol transported by any of the large, medium or small LDL subclasses (Fig. 1).

When examining triglycerides in apolipoprotein B-containing particles, *CETP* associated with lower circulating triglyceride concentrations in VLDL and IDL subclasses, while *HMGCR* had weaker effects on these measures, except in LDL subclasses (Fig. 2). *CETP* had a very strong association with higher HDL cholesterol (0.84; 95%CI: 0.76, 0.92) but *HMGCR* did not (0.04; 95%CI: -0.02, 0.10) (Fig. 3). Similarly, *CETP* was inversely associated with the total quantity of triglycerides in HDL particles (-0.23; 95%CI: -0.31,-0.15) but *HMGCR* was not (-0.03; 95%CI: - 0.09, 0.02).

**Figure 2.**
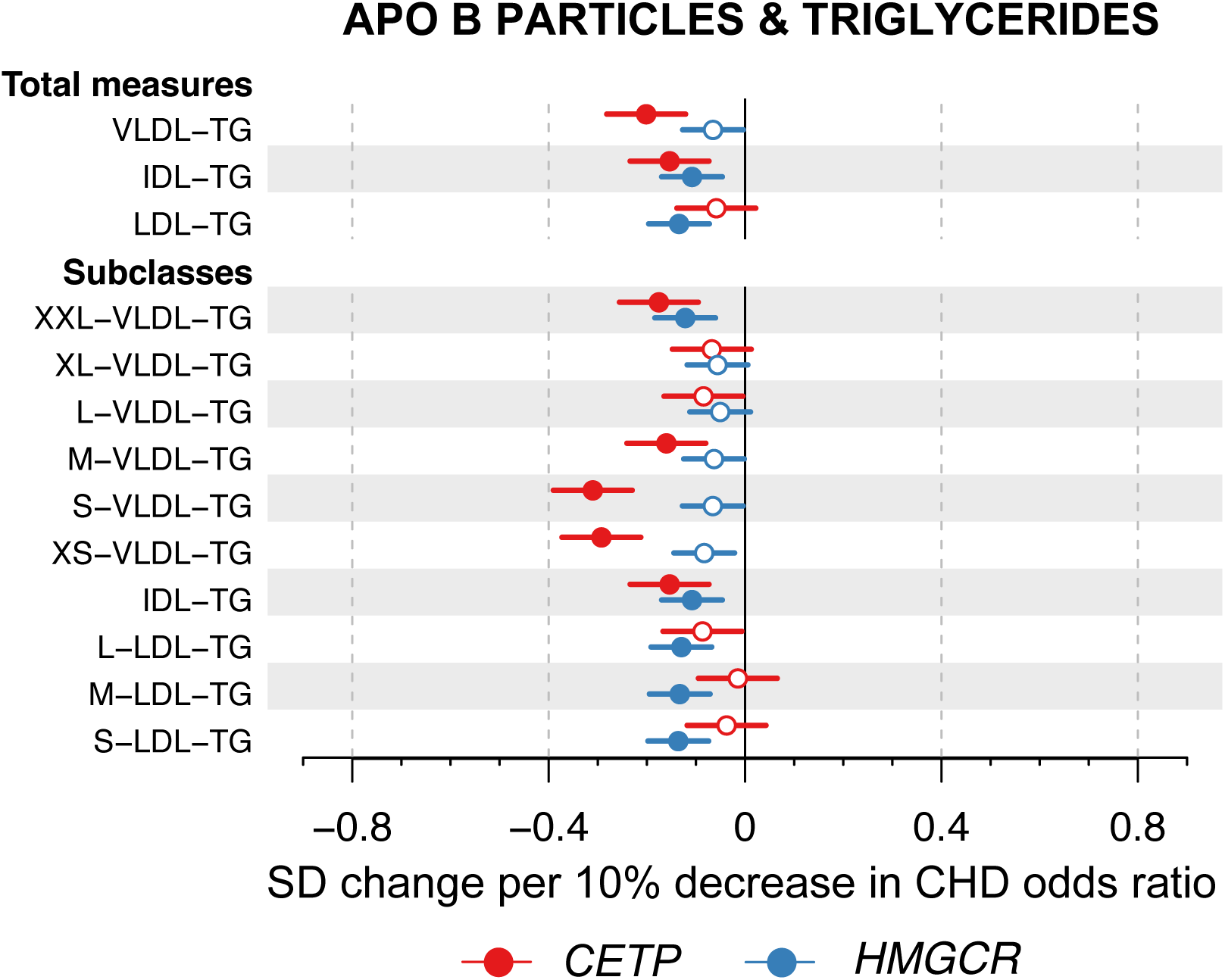
Associations of genetic variants in *CETP* rs247617 (red) and *HMGCR* rs12916 (blue) with circulating triglyceride concentrations in size-specific apolipoprotein B particles. The estimates and lipoprotein subclasses are as defined in the caption for Fig. 1.

**Figure 3.**
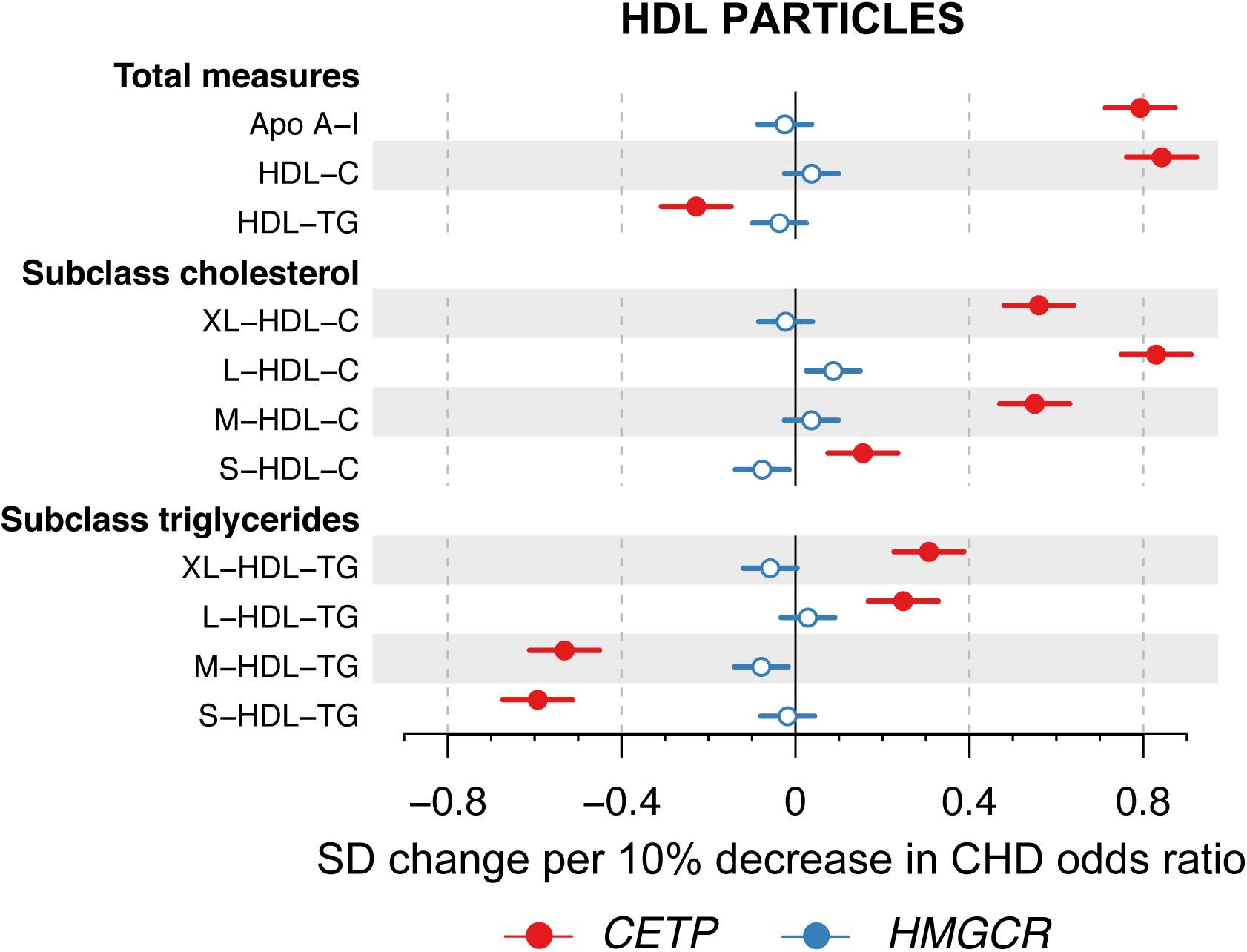
Associations of genetic variants in *CETP* rs247617 (red) and *HMGCR* rs12916 (blue) with circulating apolipoprotein A-I as well as cholesterol and triglyceride concentrations in size-specific HDL particles. The estimates are as defined in the caption for Fig. 1. The four size-specific HDL subclasses are very large (average particle diameter 14.3 nm), large (12.1 nm), medium (10.9 nm) and small HDL (8.7 nm).

The lipoprotein particle structure is biophysically constrained, generating strong correlations between lipid measures within individual lipoprotein subclasses.^25–28^ Notable differences in lipid concentrations in subclass particles would therefore suggest changes in the compositional proportions of these lipids. For genetic inhibition of CETP, the effects on circulating triglyceride concentrations in all HDL subclasses were weaker (XL-HDL and L-HDL) or even in the opposite direction (M-HDL and S-HDL) than the effects on cholesterol concentration in these subclasses (Fig. 3). Examining the genetic associations with the particle lipid compositions, the relative amount of triglycerides (in relation to all lipid molecules in the particles) was remarkably diminished in all HDL subclass particles by genetic inhibition of CETP (Fig. 4). Genetic inhibition of HMGCR did not associate with the triglyceride concentration or composition of any HDL subclass. These associations are in line with the known physiological roles of CETP and HMGCR and their inhibition.^29, 30^ In addition, as expected, *CETP* associated with higher compositions of triglycerides in most VLDL subclass particles and *HMGCR* showed directionally similar, albeit weaker associations.

**Figure 4.**
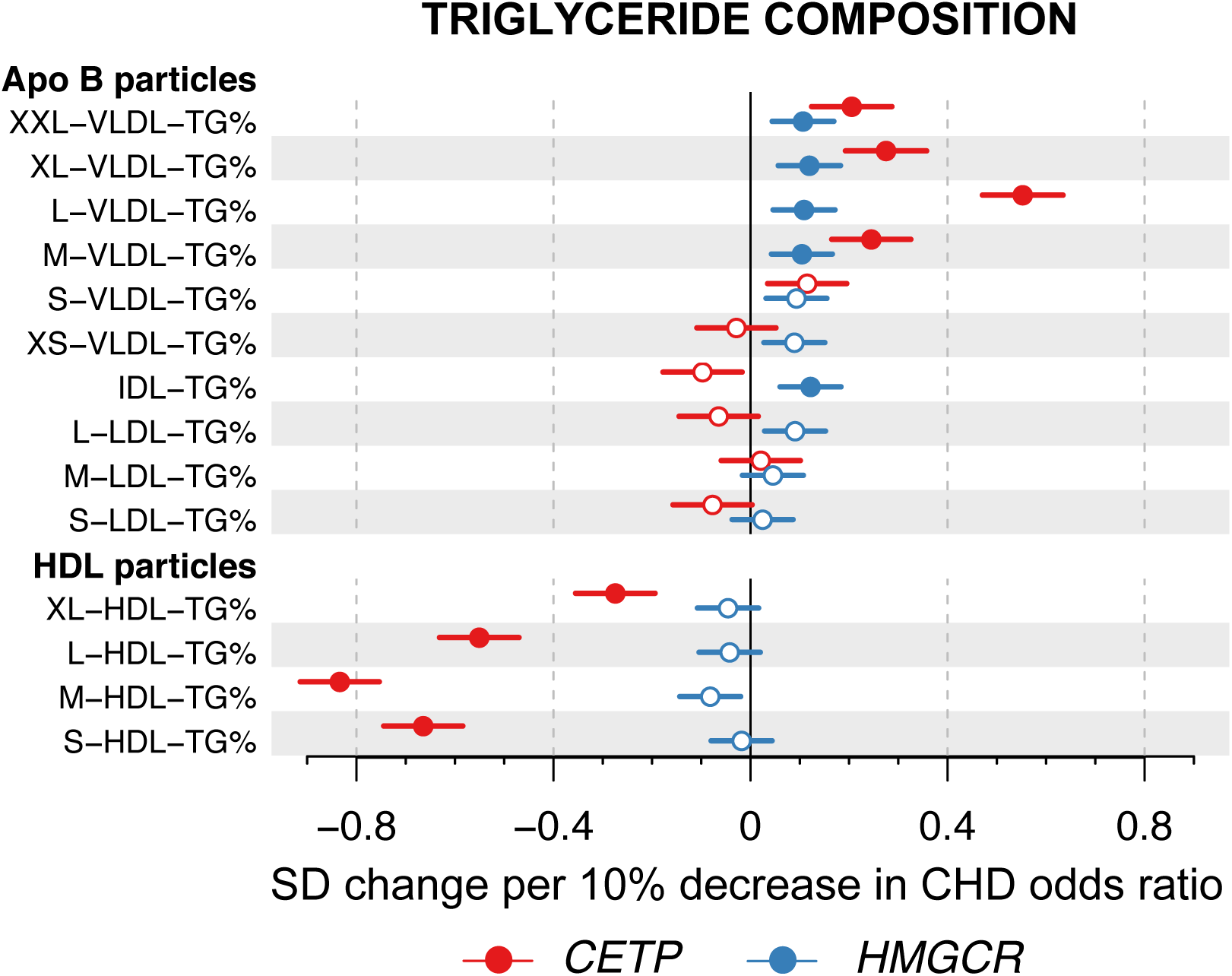
Associations of genetic variants in *CETP* rs247617 (red) and *HMGCR* rs12916 (blue) with the triglyceride composition of size-specific lipoprotein particles. The estimates are as defined in the caption for Fig. 1 and the lipoprotein subclasses are as defined in the captions for Fig. 1 and Fig. 3.

To understand the clinical relevance of these HDL-related compositional changes arising from CETP inhibition, beyond reductions in cholesterol concentrations of apolipoprotein B-containing lipoprotein particles, we studied the observational associations of lipoprotein subclass lipid concentrations and compositions with CHD in three prospective population cohorts. The triglyceride concentration of HDL was associated with incident CHD when adjusted for non-lipid cardiovascular risk factors (Fig. 5)., However, when serum cholesterol and serum triglycerides were added to the model, as expected, the associations attenuated. In contrast, the triglyceride compositions of all the HDL subclass particles were positively associated with CHD, independent of circulating concentrations of cholesterol and triglycerides, with hazard ratios around 1.3 for all HDL subclasses (Fig. 5). In addition, compositional enrichment of cholesteryl esters in the largest VLDL particles (XXL-VLDL and XL-VLDL) was associated with risk of CHD (Online Fig. 11); genetic inhibition of CETP also impacted on these traits (Online Fig. 2).

**Figure 5.**
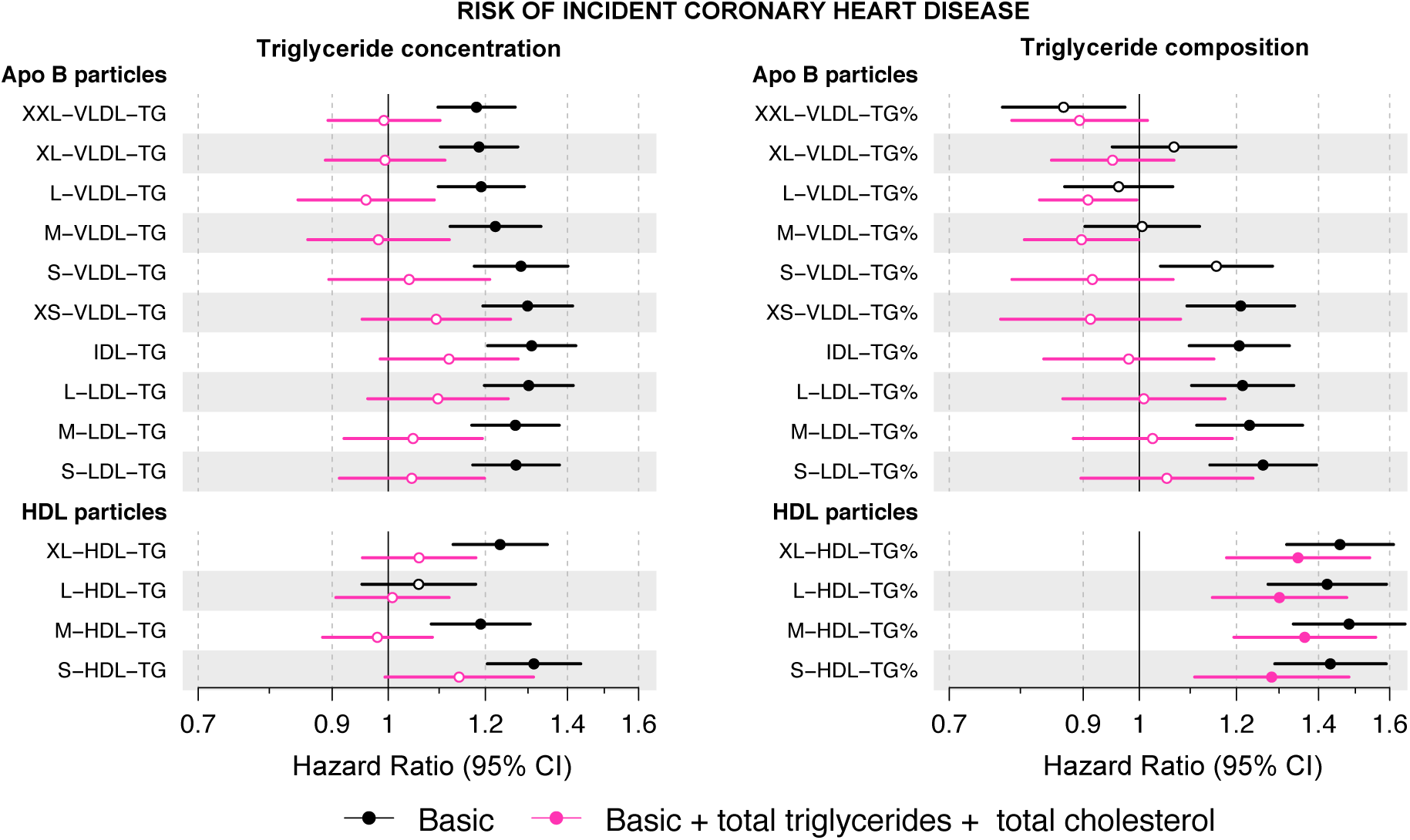
Observational associations of circulating triglyceride concentrations and triglyceride composition in lipoprotein subclass particles and risk of incident coronary heart disease. **Left pane**: Black: Hazard ratios for incident CHD per-SD higher triglyceride concentration within each size-specific lipoprotein subclass adjusted for traditional risk factors. Pink: adjusted for traditional risk factors, serum cholesterol and serum triglycerides. **Right pane**: Black: Hazard ratios for incident CHD per-SD higher percentage of triglycerides (of all lipid molecules) within each size-specific lipoprotein subclass adjusted for traditional risk factors. Pink: adjusted for traditional risk factors, serum cholesterol and serum triglycerides. Basic risk factors include age, sex, mean arterial pressure, smoking, type 2 diabetes mellitus, lipid medication, geographical region (FINRISK) and ethnicity (SABRE). Closed circles represent statistical significance of associations at P<0.002 and open circles associations that are non-significant at this threshold.

## Discussion

We used genetic variants in *CETP* and *HMGCR* to gain insight into the expected effects of therapeutic inhibition of CETP and HMG-coA reductase on circulating lipoproteins and lipids. Our data show that while *CETP* and *HMGCR* have near identical effects on Friedewald-estimated LDL cholesterol, this result masks a very different association of *CETP* and *HMGCR* with size-specific LDL cholesterol. Genetic inhibition of HMGCR showed similar effects with cholesterol across the apolipoprotein B-containing lipoproteins but genetic inhibition of CETP showed stronger associations with larger apolipoprotein B particles, namely VLDL and remnant cholesterol,^31^ but no association with cholesterol carried specifically in LDL particles defined by size.

Friedewald-estimated LDL cholesterol (and other assays such as ‘direct’ and betaquant) are non-specific measures of cholesterol.^32–34^ For example, in addition to the cholesterol in size-specific LDL particles, Friedewald LDL cholesterol also includes, to varying degrees, cholesterol in IDL, VLDL and lipoprotein(a).^35^ This non-specificity of commonly-used “LDL” cholesterol assays is under-recognized and underlies the prevailing opinion that inhibitors of HMGCR and CETP both alter LDL cholesterol. However, our data show this not to be the case: using NMR spectroscopy-based lipoprotein particle quantification, which defines individual lipoprotein subclasses based on particle size,^18, 25, 27^ our findings demonstrate that CETP has negligible effect on cholesterol in size-specific LDL particles. In this way, the use of a composite lipid measure can obscure differential associations of a therapy or gene^26^ with individual constituents of the composite, and can have clinical ramifications. For example, if a trial is powered to a given reduction in Friedewald LDL cholesterol, under the naïve assumption that the drug uniformly alters all the subcomponents, then the trial may not have the expected result if the drug has differential effects on these subcomponents. This is exemplified in the recent phase III ACCELERATE trial of evacetrapib, which was terminated for futility, and was powered to a difference in LDL cholesterol based on a composite assay.^36^ The differential effects of CETP inhibition on composite markers such as Friedewald and directly-quantified LDL cholesterol compared to apolipoprotein B concentrations identified in the subsequent phase III REVEAL trial of anacetrapib^7^ suggest that had ACCELERATE used an alternative measure of pro-atherogenic lipoproteins (e.g. apolipoprotein B or non-HDL-C^8^) to gauge the expected vascular effect, the trial may have been more appropriately powered.

This highlights the need to understand, in detail, the consequences of lipid modifying therapies on lipoproteins and lipids in order to be able to gauge whether a composite measure (such as Friedewald LDL cholesterol) can be reliably used as an indicator of the likely beneficial effect of a therapy. This is unlikely to be limited to assays for LDL cholesterol. For example, assays that quantify triglycerides, measure the summation of triglycerides across multiple lipoprotein particle categories. Drugs currently under development that target triglycerides (such as apolipoprotein C-III inhibitors^37^) have differential effects on triglycerides in lipoprotein subclass particles as demonstrated in a recent genetic study.^38^ If triglycerides within different lipoprotein subclasses have heterogeneous effects on vascular disease, a clinical trial powered to the overall concentration of circulating triglycerides may give an inaccurate portrayal of the cardiovascular consequences arising from apolipoprotein C-III inhibition.

Another key finding is that the lipid compositions of lipoprotein particles can associate with disease risk independently of total lipid concentrations. While genetic inhibition of CETP increased circulating concentrations of cholesterol in all HDL subclasses, the triglyceride composition, i.e. the percentage of triglyceride molecules of all the lipid molecules in the particle, was markedly lower in all HDL particles. Intriguingly, our observational analyses, the first to explore lipoprotein particle lipid composition with CHD outcomes, revealed that triglyceride enrichment of HDL particles associates with higher risk for future CHD, independently of total circulating cholesterol and triglycerides. The largest hazard ratio for the triglyceride enrichment in medium HDL subclass particles was of a similar magnitude (~1.3) as that for LDL cholesterol and apolipoprotein B.^39^ These findings suggest that lipoprotein particle compositions, independent of circulating lipid concentrations, could have a role in the development of CHD. While the causal role of these lipoprotein compositions remains unclear, these findings advocate the importance of moving from simple composite lipid measures towards more detailed molecular phenotyping of lipoprotein metabolism.

Key strengths of our analyses include the availability of detailed measurements of blood lipoprotein subclass concentrations and compositions from general population studies with incident CHD events, together with the availability of genome-wide genotyping. While we used *CETP* and *HMGCR* variants as genetic proxies for therapeutic inhibition, we note that the *CETP* genetic variant recapitulated the effects of CETP enzyme activity in relation to the role the enzyme has in shuttling esterified cholesterol from HDL to apolipoprotein B-containing particles in exchange for triglycerides.^29^ Furthermore, prospective population-based data of patients taking statins with blood sampling before and after the commencement of therapy showed that genetic variants in *HMGCR* robustly recapitulated the effects of statin therapy on lipoprotein subclasses and lipids.^10^

In conclusion, we have shown that, in contrast to genetic inhibition of HMG-CoA (proxying statin therapy), genetic inhibition of CETP does not alter circulating size-specific LDL cholesterol concentrations. This is masked by using conventional, non-specific assays for LDL cholesterol and may be problematic for ongoing and future clinical trials of lipid lowering therapies, especially when a non-specific marker of lipids is used to derive an expected effect of a drug with risk of disease. Our findings suggest potential additional mechanisms by which CETP inhibition could prevent CHD through reductions in the triglyceride composition of HDL particles. Our findings also call attention to the need for metabolic precision in measurements of lipoprotein lipids and in assessing the role of lipoprotein metabolism in cardiovascular disease in relation to ongoing treatment trials of novel lipid-altering therapies.

